# Secondary rewards acquire enhanced incentive motivation via increasing anticipatory activity of the lateral orbitofrontal cortex

**DOI:** 10.1101/2020.12.29.424651

**Authors:** X Yang, X Liu, Y Zeng, R Wu, W Zhao, F Xin, S Yao, KM Kendrick, RP Ebstein, B Becker

## Abstract

The motivation to strive for and consume primary rewards such as palatable food is bound by internal satiation and devaluation mechanisms, yet secondary rewards such as money may not be bound by these regulatory mechanisms. The present study therefore aimed at determining diverging devaluation trajectories for primary (chocolate milk) and secondary (money) reinforcers on the behavioral and neural level. Satiation procedures combined with a choice (Experiment 1) and an incentive delay (Experiment 2) paradigm consistently revealed decreased hedonic value for the primary reward as reflected by decreasing hedonic evaluation and choice preference, while hedonic value and preferences for the secondary reward increased. Concomitantly acquired functional near-infrared spectroscopy (fNIRS) data during the incentive delay paradigm revealed that increasing value of the secondary reward was accompanied by increasing anticipatory activation in the lateral orbitofrontal cortex, while during the consummatory phase the secondary reinforcer associated with higher medial orbitofrontal activity irrespective of devaluation stage. Overall, the findings suggest that – in contrast to primary reinforcers - secondary reinforcers can acquire progressively enhanced incentive motivation with repeated receipt, suggesting a mechanism which could promote escalating striving to obtain secondary rewards.

## 1. Introduction

A salient survival function of the brain is to direct behavior towards primary rewards, such as food, shelter and mates. Behavioral control mechanisms regulate subjective reward values by taking into account both the external (environment) as well as the internal state of the organism and thus facilitate goal-directed behaviors [1]. Successful goal-directed decision making necessitates a dynamic balance and integration of information from the external environment and the current internal state to maximize resource allocation while minimizing energy expenditure. With respect to the dynamic representation of experienced reward value, this process thus relies on external variables such as the overall rewarding properties of the stimuli (e.g. palatability or calories of food) as well as the internal milieu including the organism’s homeostatic (i.e. thirst or hunger) and hedonic state [2, 3].

Reward evaluation and actual food consumption are strongly influenced by internal physiological factors that signal the homeostatic state of the organism, such as glucose, insulin, or leptin blood level [4–6], and are dynamically modulated as optimal levels of homeostasis are approached. Furthermore, given that the organism’s hedonic valence contributes to reward evaluation and subsequent intake of food [7, 8], eating behavior hence is partially based on its subjective hedonic valence. Additionally, hedonic valence of food depends on the organism’s previous experience with the food. Intriguingly, the consumption of any specific food leads to temporary sensory-specific satiety, viz. its devaluation is accompanied by the consequent renewal in appetite resulting from the exposure to a new food [9–13]. In conclusion, the consumption of natural reinforcers such as palatable food is regulated by adaptations in the homeostatic and hedonic state of the organism that mediate consummatory behavior via a process of devaluating its reinforcing properties.

Human, as well as animal, behavior is not only driven by innate natural (primary) rewards but also by learned (secondary) rewards which are not inherently rewarding or survival relevant but rather gain their subjective motivational value via learned associations with primary rewards [14]. In particular, among the multitude of stimuli that can acquire motivational significance from chocolate cake to cigarettes via associative learning mechanisms, money conspicuously stands out and has enormous motivational valence for many people. This is not surprising since from ancient times money is the tender that enables an individual to most easily obtain primary rewards. It is worth considering that in comparison to primary rewards, secondary rewards are not necessarily bound by an innate homeostatic balance. For example, use of drugs of abuse such as opioids is often characterized by escalating amount of use likely based on dysregulated reward processing that leads to dysregulated goal-directed behavior [15, 16]. Similarly, money can become an end in itself and its pursuit is often unconstrained by normal hedonic valence or homeostatic balance leading to dysregulated and self-destructive behavior, viz. King Midas who according to <u>Aristotle</u>, legend held that Midas died of starvation as a result of his “vain prayer” for the gold touch. A more recent example is the figure of Jordan Belfort, a stockbroker involved in fraud and other nefarious activities and depicted in the movie ‘The Wolf of Wall Street’.

Animal models and neuroimaging findings in humans suggest that primary and secondary reward types are processed in partly overlapping neural systems, including core regions of the meso-limbic dopamine circuit such as the nucleus accumbens and amygdala, as well as the prefrontal cortex, particularly the orbitofrontal cortex (OFC) [14, 17–19]. To date, most functional magnetic resonance imaging (fMRI) studies have focused on the neural reactivity to either primary rewards such as highly palatable food, or secondary rewards such as money but recent studies have begun to directly compare neural responses to different reward categories in humans[18–20]. These studies revealed evidence that primary rewards such as palatable food or erotic stimuli induce a stronger engagement of the anterior insula, the amygdala and posterior lateral orbitofrontal cortex (OFC) regions whereas monetary-rewards (secondary) are specifically encoded in the most anterior portion of the OFC [17–19] suggesting that secondary rewards such as money are represented in more recently evolved brain regions. The OFC not only plays an important role in the initial evaluation of rewards but also critically mediates adaptive changes in reward evaluation, including hedonic devaluation represented by sensory-specific satiety [7, 18, 21–23].

A growing consensus converges on the concept that the *lateral* OFC (lOFC/vlPFC)[24] likely represents the agent’s initial position (initial state) regarding assignment of value within that task map and hence determines which actions are available as a consequence of the initial state. In contrast the medial OFC, often referred to as the ventromedial prefrontal cortex (mOFC/vmPFC) in humans, represents the agent’s future state and action outcomes with respect to general value coding within the task map. This in turn partially determines which actions are selected to achieve desired action outcomes [11, 23, 25–27]. In humans an increasing number of studies employed fMRI to examine satiety-related devaluation of primary reinforcers such as palatable food. These studies demonstrated that the lateral posterior OFC activity was modulated after food devaluation, whereas the vmPFC/mOFC was sensitive to the general value of reward. Moreover, the lateral and medial prefrontal systems interact such that the vmPFC/mOFC integrates information from the vlPFC/lOFC to guide goal-directed behavior[11, 28].

Despite an increasing number of human fMRI studies that investigated the neural devaluation of rewarding food/odor stimuli in different metabolic states (hunger vs. satiety) they were often confounded by methodological limitations inherent to fMRI. These limitations include high motion sensitivity that interferes with satiety protocols implemented during data acquisition and thus neural activation is commonly examined before and after satiety-induction [11, 20, 29]. Notably, a direct comparison of the behavioral and neural processes of devaluation between primary and secondary rewards has yet to be conducted.

Against this background the present study incorporated two experiments that aimed at comparing reward devaluation implemented using a satiation procedure of primary (chocolate milk) and secondary (money) rewards both on the behavioral (experiment 1) and neural (experiment 2) level. To facilitate a sensitive assessment of the behavioral and neural differences, the study encompassed two experiments in independent samples. Experiment 1 employed a forced-choice paradigm in N = 30 healthy male participants to determine differences on the behavior level, whereas experiment 2 combined an incentive delay paradigm with concomitant neural assessments of the prefrontal activation in N = 35 healthy male participants. For the neural assessments, functional near-infrared spectroscopy (fNIRS) was employed. fNIRS is a noninvasive optical imaging technique that indirectly assesses neural activity by measuring changes in oxygenated and deoxygenated hemoglobin in tissues using near-infrared light. Despite limitations, viz. the technique is restricted by penetration depth and spatial resolution in comparison to fMRI, fNIRS has nevertheless gained considerable traction in cognitive and clinical neuroscience. In comparison to fMRI, fNIRS has the advantage of a lower sensitivity to motion artifacts and lower susceptibility to interferences in prefrontal regions. Finally, fNIRS can be applied during naturalistic settings [30–34], which makes it highly attractive for monitoring neural changes over the course of reward consummation and devaluation procedures.

Based on the sensory-specific satiety concept [9–13], we hypothesized an increasing devaluation of the primary reward over the satiation procedure, while we expected that the motivational significance and hedonic value of the secondary reward would increase. Given the important contribution of the OFC in reward and reward devaluation as well as the functional heterogeneity of the OFC [35, 36], we predicted that (1) rewarding properties of monetary reward would be reflected by higher activation in the medial OFC due to the role of this region in coding the general reward value that guides goal directed behavior, while (2) the increase in hedonic value of the secondary reward would be accompanied by increasing activation of lateral subregions of the OFC due to its role in representing a reinforcer’s momentary (dynamic) position/state in which it calculates the current value of rewards.

## 2. Methods and materials

### 2.1 Participants

The present study employed a satiation procedure to determine differential devaluation of primary (natural) and secondary (learned) rewards both on the behavioral and neural level. To facilitate a sensitive assessment of the behavioral and neural differences, the study encompassed two experiments carried out in independent samples. Experiment 1 used a forced-choice task paradigm in N=30 healthy male participants (M age = 22.17 ± 2.04 years, range = 18-26; M BMI = 21.19 ± 1.56, range = 18.61 − 23.24) to determine preference differences on the behavioral level. Experiment 2 combined an incentive delay paradigm with fNIRS in N=35 healthy male participants (M age = 21.74 ± 2.02 years, range = 18 − 25, M BMI = 20.91 ± 1.52, range = 18.38 − 23.67) to determine neural differences in the devaluation of natural and learned rewards. Inclusion criteria for both experiments included right handedness, male sex, and age > 18 years. To ensure an overall high rewarding property of the natural reward in the present sample only subjects who reported that they liked chocolate milk during the pre-screening were invited (likeability rating > 7 on a 10-point rating scale). The following exclusion criteria were applied: BMI < 18.5 or > 23.9, current dietary food restrictions, no lifetime diagnosis of psychiatric or neurological disorders, including eating disorders, and no current or regular use of psychotropic substances, including nicotine. Experimental procedures adhered to the latest revision of the Helsinki Declaration and had approval by the local ethics board. All subjects were recruited via advertisements, provided written informed consent and received monetary compensation for participation.

### 2.2 Experimental procedures

Participants were instructed to abstain from food for at least 6h before the experiment. To account for circadian variations of hunger and appetite all experiments were scheduled in the early evening (start of the experiments between 5-6PM, after regular lunch between 11-12AM). Prior to the experimental session, participants were offered water to control for unspecific effects of thirst and underwent a practice session of the paradigms to validate comprehension of the experimental instructions. Before and after the experiments blood glucose levels (measured by OneTouch Ultra Easy blood glucose meter, Johnson & Johnson, New Jersey, United States, a procedure validated by [37, 38]) as well as subjective ratings of the desire to eat, fullness, satiety and pleasantness ratings of each stimulus category (chocolate milk, money, water were assessed via visual analogue scales (VAS).

#### Experiment 1 (behavioral level) – choice reward paradigm

Experiment 1 employed a choice reward outcome paradigm. During this computerized paradigm, participants were presented with 90 trials. On every trial participants had to choose between receiving one of two simultaneously presented rewarding outcomes. Choice options included (a) a primary reward (chocolate milk, 20 ml), (b) a secondary reward (money, 0.1 RMB (0.015 $ US), or (c) a low primary reward (20 ml water) option which additionally served as a control condition to account for unspecific devaluation of the sensory properties of liquid. Specifically, the following combination of options were displayed: chocolate milk vs. money, chocolate milk vs. water, and money vs. water. 30 trials per combination were displayed in a pseudorandomized and counterbalanced order. Each trial started with a 3s ‘choice’ displaying the option and participants indicated their choice of one of the rewards by button press (left and right index finger, position of the options counterbalanced). Immediately after the ‘choice’ phase participants received and consumed the chosen reward (chocolate milk, money or water; duration, 8s). Pleasantness ratings for the three rewards were collected following every 5^th^ trial using a 0 − 9 Likert rating scale (9 corresponding to highest pleasantness) (**Figure 1A**).

**Figure 1:**
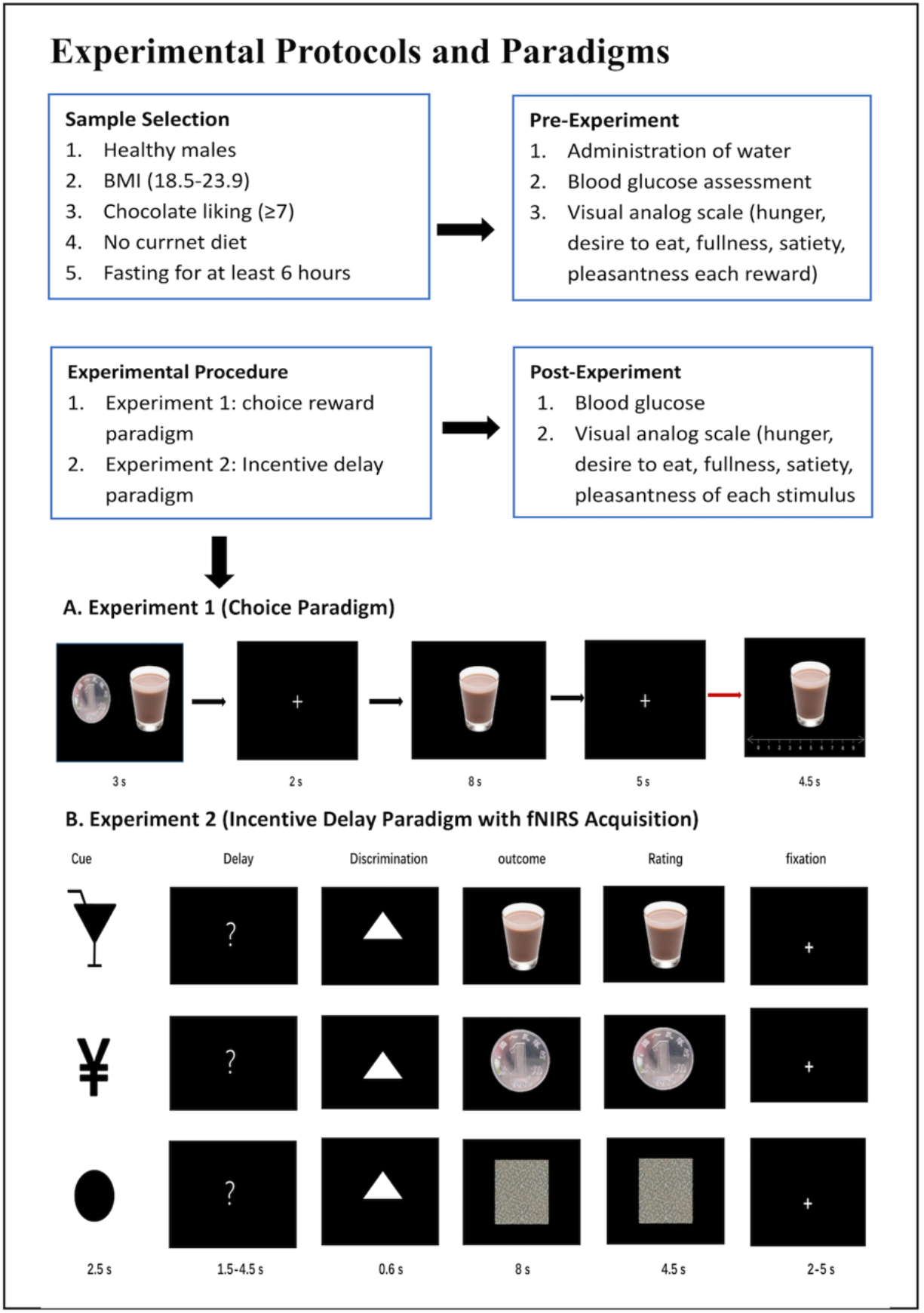
Experimental protocols and paradigms. **(A**) Experiment 1 employed a choice reward outcome paradigm. Each trial started with a 3s ‘choice’ period displaying the option and participants chose one of the presented reward options by button press (left and right index finger, position of the options counterbalanced). Subsequently participants received and consumed the chosen reward (chocolate milk, money or water; duration, 8s). Pleasantness ratings for the three reward options were collected following every 5 trials using a 0-9 rating scale (9 corresponding to highest pleasantness). **(B)** Experiment 2 combined an Incentive delay paradigm with fNIRS acquisition: The trial incorporated separate phases to assess reward expectation and outcome. During a trial subjects were first presented a cue informing them about the type of reward. Three cues were presented: chocolate milk, money, and nothing which served as control condition. After a short delay and a target discrimination task, subject received the outcome. Reward outcomes consisted of chocolate milk (top) or money (middle) or nothinng respectively and were followed by a pleasantness rating. Non-reward and control trials displayed a scrambled picture as outcome (bottom).

#### Experiment 2 with concomitant fNIRS assessment – incentive delay paradigm

Experiment 2 aimed at determining the neural basis of reward-type dependent devaluation by concomitant fNIRS assessment during a modified version of a previously validated incentive delay paradigm [18]. The paradigm included two successive runs each consisting of 60 trials per run (**Figure 1B**). Experimental trials were designed to allow examination of reward anticipation and outcome. Trials were separated by a jittered inter-trial interval and included a cue, delay phase (reward anticipation), discrimination task, outcome phase (reward outcome) and pleasantness rating. The cue signaled the potential reward of the trial, signaling that either a primary reward (chocolate milk), secondary reward (money) or nothing (control) could be gained during the present trial. The ‘nothing’ cue indicated that no reward could be gained during the trial and served as control condition. Following a variable delay period (1.5 − 4.5s) subjects performed a discrimination task during which they had to respond correctly to a target shape within 0.6s. The target shape was drawn at random on each trial and could be either a triangle (‘2’ button press) or a square (‘3’ button press). Subjects were told that a correct response would lead to the signaled outcome (chocolate milk, 20 ml; money, 0.1 RBM or nothing in the control condition). Erroneous or too slow responses led to no reward. However, in order to control for the number of trials per outcome condition in the fNIRS analysis, the correct response rate was predetermined (80% correct rate). Following the outcome phase during which subjects received the reward (either drinking the chocolate milk or receiving the money), subjective pleasantness ratings were acquired using a 0-9 Likert rating scale (with 0 corresponding to no pleasure and 9 corresponding to the highest pleasure experience). Following non-rewarded trials and the control condition a scrambled picture was presented.

#### Experiment 2 - fNIRS data acquisition and preprocessing

Given the pivotal role of the prefrontal cortex (PFC), particularly the OFC, in reward processing including selective food devaluation [39, 40], fNIRS acquisition focused on the prefrontal cortex (PFC). Hemodynamic response (HR) signals in the PFC were collected using a NIRSport fNIRS system (NIRx Medical Technologies LLC, Minneapolis, MN, USA) with 8 sources and 8 detectors operating at two wavelengths (760 and 850 nm) with a sampling rate of 8.93 Hz. An optode-set of 7 sources and 7 detectors was used leading to 19 source-detector pairs. Optodes were placed with a source-detector distance of 30mm and positioned according to the International 10-20 system [41] and previous fNIRS studies examining the PFC [42, 43]. In line with these previous studies the channels were allocated to the ventrolateral prefrontal cortex (vlPFC/lOFC, channels: 1-4, right vlPFC/lOFC and channels:16-19, left vlPFC/lOFC) and the ventromedial prefrontal cortex (vmPFC/mOFC, channels 5-15). Oxygenated (HbO) and deoxygenated (HbR) signals were assessed. Consistent with previous studies employing fNIRS to examine reward-related PFC activation [44, 45], and also gain evidence for higher signal-to-noise ratio as well as amplitude of the HbO relative to the HbR during task engagement [46, 47], the primary analysis focused on the HbO signal.

FNIRS data processing and analysis procedures were accomplished using SPM-fNIRS [48], which is based on SPM12 [49]implemented in MATLAB (Mathworks, Natick, MA). Raw intensity data were converted to hemoglobin changes using the modified Beer-Lambert Law [50]. For spatial pre-processing the fNIRS, channels were registered to the native space of each participant. Temporal pre-processing included the removal of physiological noise using filtering (band-stop filter: 0.12-0.35 – 0.7-2 Hz). In order to reduce motion artifacts a previously validated approach based on moving standard deviation and spline interpolation [51] with a 3000ms moving window length was employed. The threshold factor for motion detection was set to 3 and for smoothing and factor-motion artifact was set to 5. Next low-frequency confounds were removed by applying a high-pass filter based on a discrete cosine transform (DCT-cut-off period :128), and high frequency noise was reduced using a low-pass Gaussian filter [Gaussian smoothing with full width at half maximum (4 s)] [52].

A generalized linear model (GLM) approach was employed to model the task-related hemodynamic response on the individual level, including regressors for the cue, delay, discrimination, outcome rating and fixation period. The design matrix included condition-specific regressors for reward anticipation (anticipation of chocolate milk, money, control) and outcome (chocolate milk, money, control). The incentive delay task was usually used to investigate ‘wanting’(desire to reward) and ‘liking’(pleasure during receipt of rewards)[20] .The first-level design matrix was convolved with a standard HR function as provided in the SPM-fNIRS package. For the group level (second level) analysis channel-specific beta values were obtained from each subject. To control for individual differences in baseline neural activation and unspecific neural activation related to, e.g. basal visual and attention processes, the control condition (nothing) was subtracted from each reward condition (money, chocolate milk). The channel- and condition-specific beta-estimate contrasts were subsequently subjected to a group-level analysis by means of a repeated measures ANOVA with the factors reward condition (2: chocolate milk, money), time (2: run 1, run 2) and channel (1-19) and subsequent False Discovery Rate (FDR) corrected post-hoc tests [53].

## 3. Results

One subject reported having eaten during the 6h period before experiment 1 and was excluded, leaving n = 29 subjects for the analysis. For experiment 2, n = 3 participants had to be excluded due to an unusual high error rate during the experiment (z-values >3) leaving n = 32 participants for data analysis. The pre- and post-experiment ratings as well as blood glucose levels, revealed that at the start of both experiments, subjects were in a hungry state and desired to eat, viz. they were not satiated or full. Examination of changes across the experimental session by means of paired t-tests further established that these ratings indeed significantly changed in the expected direction (**Table 1**).

**Table 1:**
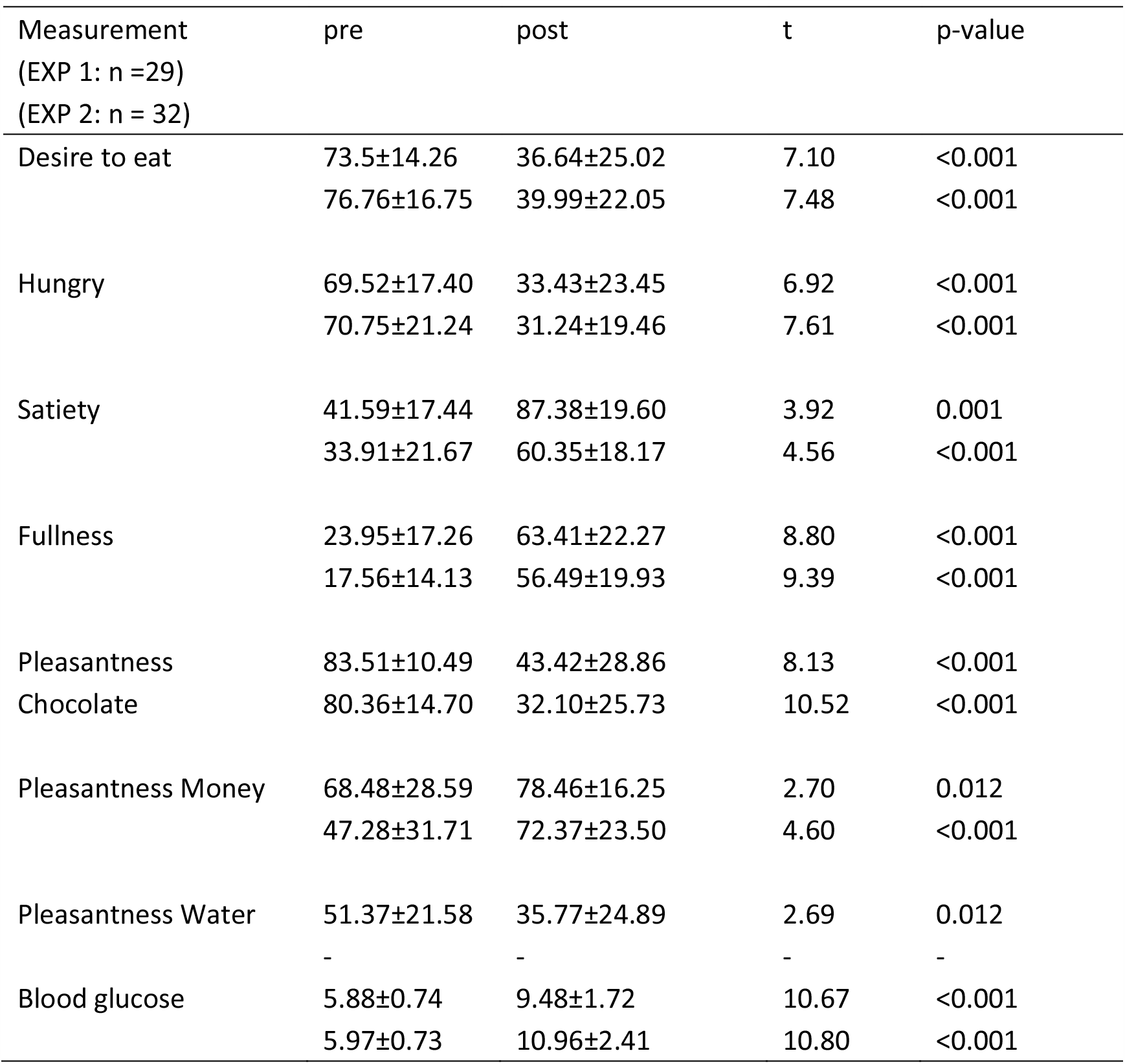
VAS (range, 0-100) and blood glucose immediately pre- and post-test(mean±sem) Experiment 1 (EXP 1) & Experiment 2 (EXP 2)

### 3.1 Changes in pleasantness ratings before and after the experiment

Changes in pleasantness ratings before and after the experiment (via visual analogue scales [VAS]) were examined by means of repeated measures ANOVAs with timepoint (pre-, post-experiment) and reward condition as factors. For both experiments significant main effects of timepoint and interaction effects were observed (both Ps < 0.001), with post-hoc paired t-tests indicating that across both experiments the pleasantness for chocolate milk and water in experiment 1 decreased, whereas the pleasantness for money increased in the two experiments (**Figure 2A & B**). Together these findings confirm sensory specific devaluation for the primary reward but the opposite pattern for money (secondary reward) was observed.

**Figure 2.**
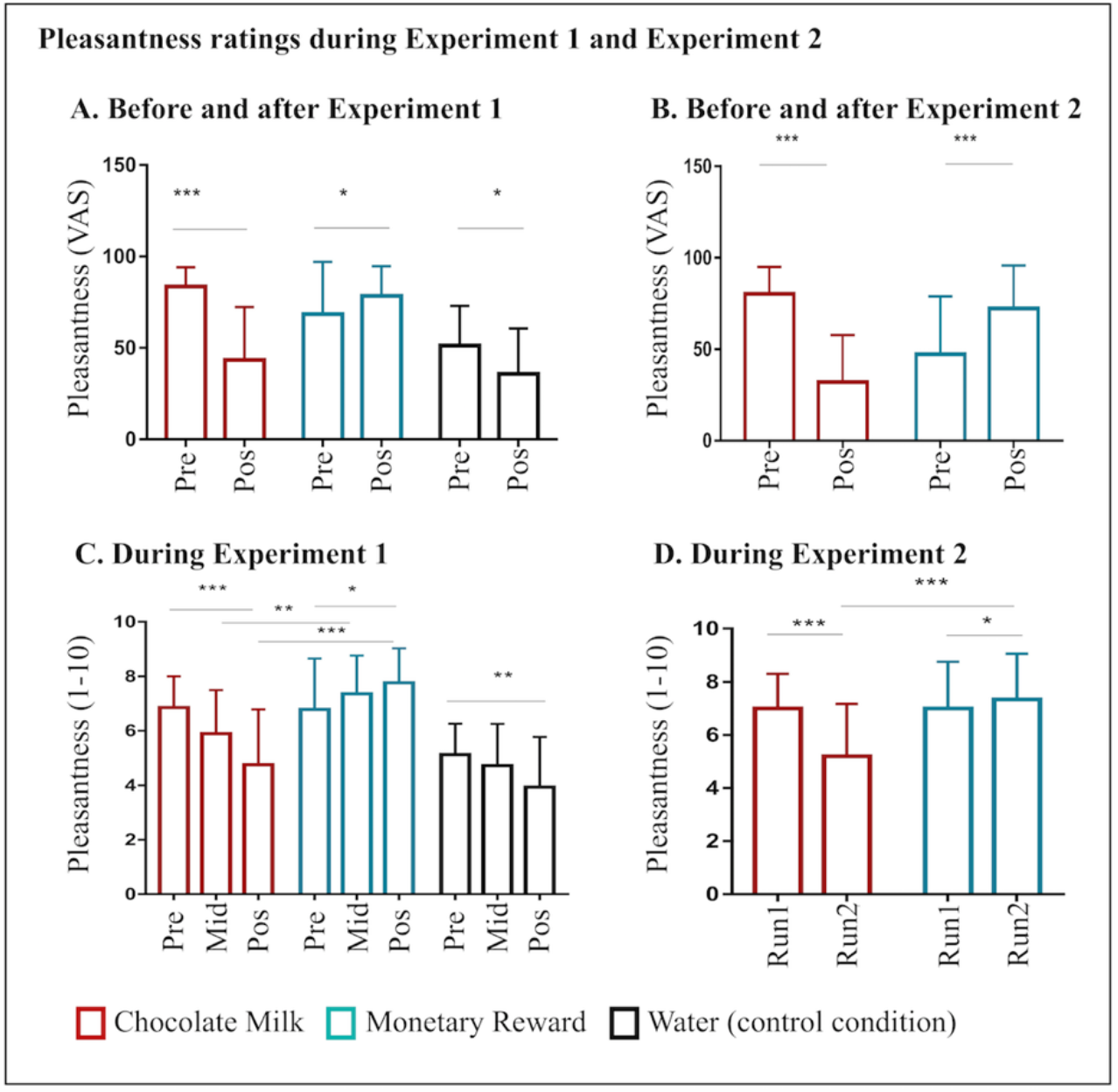
Pleasantness ratings for Experiment 1 and Experiment 2. Pleasantness ratings before and after experiment 1 (**A**) and experiment 2 (**B**) as assessed via Visual Analog Scales (VAS, 0-100). Pleasantness ratings during experiment 1 (**C**) and experiment 2 (**D**) acquired during the experimental paradigms. Pleasantness ratings were assessed for chocolate milk, money (both experiments) and water (experiment 1). Pleasantness ratings for chocolate milk and water decreased over the experiments, whereas the pleasantness for money increased. Abbreviations; pre, pre-experiment assessment; pos, post-experiment assessment; mid, mid-experiment assessment; run1, first run of the incentive delay paradigm; run2, second run of the incentive delay paradigm *P _(Bonferroni corrected)_ < 0.05, ** P _(Bonferroni corrected)_ < 0.01, *** P _(Bonferroni corrected)_ <0.001.

### 3.2 Pleasantness ratings over the course of experiment 1 and 2

#### Experiment 1

For the analysis of changes in pleasantness experience over the course of the experiment pleasantness ratings reward-specific bins including 6 condition-specific pleasantness trials were computed (mean values of the early, 1-6, mid: 7-12, late: 13-18 trials in experiment 1, with a total of 90 trials, pleasantness ratings for the three rewards were collected following every 5^th^ trial, totaling 18 ratings for each reward type) and subjected to a corresponding repeated measures ANOVA. Examination of the pleasantness ratings by means of a repeated measures ANOVA with the factors including reward type condition (money, chocolate, water) and timepoint (early, mid, late), revealed significant main effects of condition (F(2, 56) = 41.46, P < 0.001, partial η^2^ = 0.60) and timepoint (F(2, 56) = 13.02, P < 0.001, partial η^2^ = 0.32) as well as a significant interaction effect between the factors (F(4, 112) = 19.32, P < 0.001, partial η^2^ = 0.41). Post-hoc t-tests showed that reward ratings for both primary rewards (chocolate milk, water) decreased whereas ratings for the monetary reward increased (chocolate milk_pre vs. chocolate milk_post, P _(Bonferroni corrected)_ < 0.001, Cohen’s d = 1.27; water_pre vs. water_post, P _(Bonferroni corrected)_ < 0.01, Cohen’s d = 0.78; money_pre vs. money_post, P _(Bonferroni corrected)_ < 0.05, Cohen’s d = 0.62). Directly comparing the timepoint-specific ratings further revealed that chocolate milk and money did not significantly differ during the early phase (chocolate milk_pre vs. money_pre, p = 0.85) and that both were rated as more pleasant than water (chocolate milk_pre vs. water_pre, P _(Bonferroni corrected)_ < 0.001, Cohen’s d = 1.51; money_pre vs. water_pre, P _(Bonferroni corrected)_ < 0.001, Cohen’s d = 1.07). Intriguingly, money had significantly higher ratings compared to chocolate during mid and late phases of the experiment (chocolate milk_mid vs. money_mid, P _(Bonferroni corrected)_ < 0.01, Cohen’s d = 0.97; chocolate milk_post vs. money_post, P _(Bonferroni corrected)_ < 0.001, Cohen’s d = 1.78) with both continuing to receive higher rating scores than water (chocolate milk_mid vs. water_mid, P _(Bonferroni corrected)_ < 0.01, Cohen’s d = 0.75; money_mid vs. water_mid, P _(Bonferroni corrected)_ < 0.001, Cohen’s d = 1.79), albeit only money had significantly higher pleasantness ratings compared to water in the late phase of the experiment (money_late vs. water_late, P _(Bonferroni corrected)_ < 0.001, Cohen’s d = 2.43). Importantly, both the interaction and post hoc tests remained significant when unspecific effects of devaluation of fluids on the pleasantness experience of chocolate milk were accounted for (chocolate > water) and compared to pleasantness ratings during the money condition. Specifically this analysis revealed a main effect of condition (F (1, 28) = 292.30, P < 0.001, partial η^2^ = 0.91) as well as a significant interaction effect (F (2, 56) = 10.09, P < 0.001, partial η^2^ = 0.27). Post hoc tests confirmed a decrease of the rewarding effects for the corrected chocolate ratings P _(Bonferroni corrected)_ < 0.05, Cohen’s d = 0.53). (**Figure 2C**).

### Experiment 2

For experiment 2, a repeated measures ANOVA with reward condition (money, chocolate) and timepoint (run1, run2) revealed significant main effects of condition (F(1, 31) = 8.58, P < 0.01, partial η^2^ = 0.22) and time (F(1, 31) = 18.17, P < 0.001, partial η^2^ = 0.37) as well a significant interaction effect (F(1, 31) = 56.52, P < 0.001 partial η^2^ = 0.65). Post-hoc t-tests for the interaction effect revealed that the pleasantness ratings for primary reward (chocolate milk) decreased (run 1 vs. run 2, P _(Bonferroni corrected)_ < 0.001, Cohen’s d = 1.08) whereas those for secondary reward (money) increased (run 1 vs. run 2, P _(Bonferroni corrected)_ < 0.05, Cohen’s d = 0.20). Directly comparing the ratings during the two runs further revealed that the reward conditions did not significantly differ during run 1 (P = 0.99, NS). However, money had significantly higher ratings compared to chocolate during run 2, P _(Bonferroni corrected)_ < 0.001, Cohen’s d = 1.16) (**Figure 2D**).

Summarizing, across both experiments, the pleasantness ratings for chocolate milk provided clear evidence for sensory-specific satiety/devaluation whereas the motivational significance of money increased.

### 3.3 Behavioral results experiment 1

For the choice reward outcome paradigm, the number of each type of reward chosen, and RTs were examined using repeated measures ANOVAs. The dependent variables were the number of each reward chosen, the reaction times, reward condition (chocolate milk vs. money vs. water) and timepoint (early, mid, late phase of the experiment). Consistent with the assessment of the pleasantness ratings, changes over the course of the experiment were examined using time bins (bin 1: trials 1-30, early; bin 2: trials 31-60, mid; bin 3: trials 61-90, late). Given that some subjects never chose water, the control condition was not included in the RT analysis.

Examining the number of each type of reward chosen revealed a significant main effect of condition (F (2, 56) = 92.04, P < 0.001, partial η^2^ = 0.77) and a condition x timepoint interaction effect (F (4, 112) = 19.04, P < 0.001, partial η^2^ = 0.41), suggesting that the value of chocolate and money changed differentially over the experiment. Furthermore, the interaction effect following post-hoc t-test revealed that the number of choices for chocolate milk decreased over the experiment, whereas the number of choices for money increased over the experiment (chocolate milk_pre vs. chocolate milk_pos, P _(Bonferroni corrected)_ < 0.001, Cohen’s d = 1.14; money_pre vs. money_pos, P _(Bonferroni corrected)_ < 0.001, Cohen’s d = 1.23). The number of water choices did not change over the experiment (P = 0.49, NS). Directly comparing the reward types within each time bin further revealed that at bin 1 choices of chocolate milk and money were comparable (P = 0.16, NS), and both were chosen more often than water (chocolate milk_pre vs. water_pre, P _(Bonferroni corrected)_ < 0.001, Cohen’s d = 2.25; money_pre vs. water_pre, P _(Bonferroni corrected)_ < 0.001, Cohen’s d = 2.77). At bin 2 chocolate milk was chosen significantly less often than money (chocolate milk_mid vs. money_mid, P _(Bonferroni corrected)_ < 0.001,, Cohen’s d = 2.15), but both continued to be chosen more often than water (chocolate milk_mid vs. water_mid, P _(Bonferroni corrected)_ < 0.01, Cohen’s d = 1.20; money_mid vs. water_mid, P _(Bonferroni corrected)_ < 0.001, Cohen’s d = 4.27), at bin 3 chocolate milk was chosen significantly less often than money (chocolate milk_post vs. money_post, P _(Bonferroni corrected)_ < 0.001, Cohen’s d = 3.79) and both continued to be chosen more often than water (chocolate milk_post vs. water_post, P _(Bonferroni corrected)_ < 0.05, Cohen’s d = 1.03; money_post vs. water_post, P _(Bonferroni corrected)_ < 0.001, Cohen’s d = 6.03)(**Figure 3A**)

**Figure 3.**
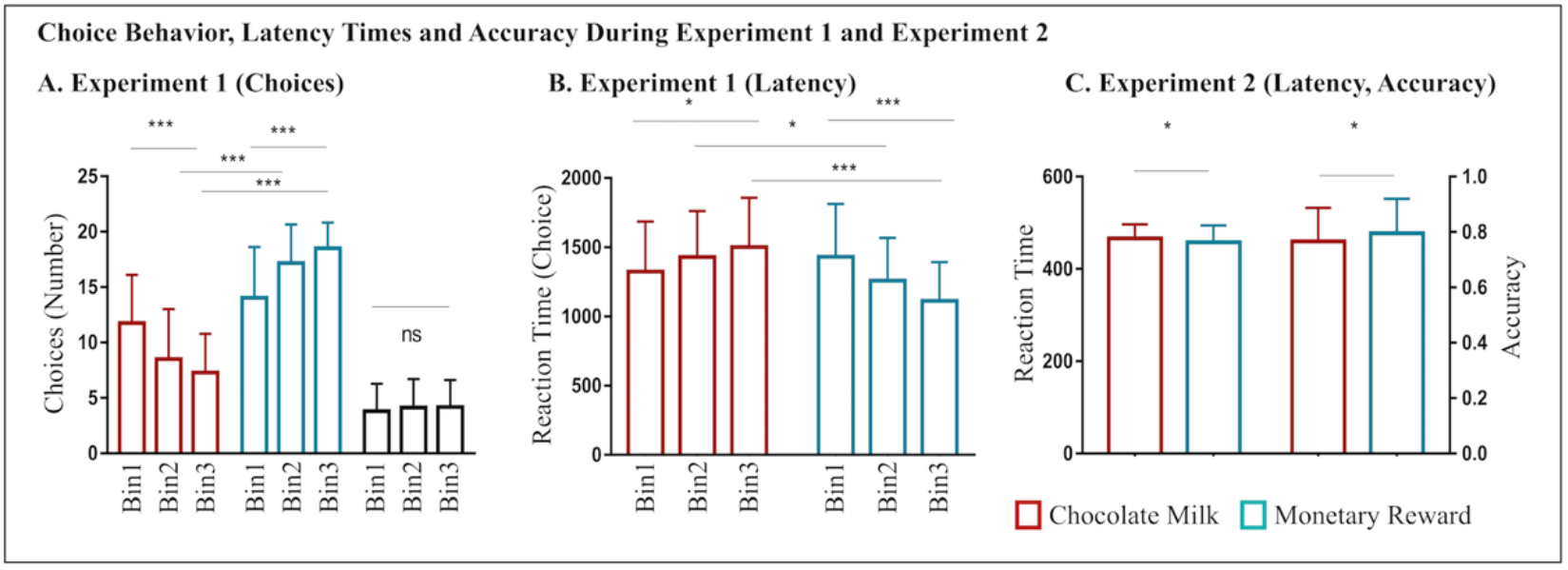
Choice Behavior, Latency Times and Accuracy During Experiment 1 and Experiment 2. **(A)** The number of choices for chocolate milk decreased over the experiment, whereas the number of choices for money increased over the experiment. **(B)** Reaction times for chocolate milk choices increased, while reaction times for money choices decreased. **(C)** In experiment 2 money had generally higher accuracy and faster responses than chocolate *P _(Bonferroni corrected)_ < 0.05, ** P _(Bonferroni corrected)_ < 0.01, *** P _(Bonferroni corrected)_ <0.001.

Examination of RTs revealed a significant main effect of reward type (F (1, 28) = 9.36, P < 0.01, partial η^2^ = 0.25) and a reward type × timepoint interaction effect (F (2, 56) = 36.75, P < 0.001, partial η^2^ = 0.57). Post-hoc t-tests revealed that RTs for chocolate milk choices increased, whereas RT of money choices decreased (chocolate milk_pre vs. chocolate milk_post P _(Bonferroni corrected)_ < 0.05, Cohen’s d = 0.49; and money_pre vs. money_pos, P _(Bonferroni corrected)_ < 0.001, Cohen’s d = 0.95). Directly comparing the two rewards during each bin revealed that there was no significant difference between money and chocolate milk during the bin 1 time period (P = 0.04), whereas RTs for money were significantly faster than for chocolate milk during bins 2 and 3 (chocolate milk_mid vs. money_mid, P _(Bonferroni corrected)_ < 0.05, Cohen’s d = 0.53; and chocolate milk_post vs. money_post, P _(Bonferroni corrected)_ < 0.001, Cohen’s d = 1.21) (**Figure 3B**).

### 3.4 Behavioral results experiment 2

Both the incentive delay task accuracy (ACC) and RTs in the discrimination task were analyzed using repeated measures ANOVAs including the factors reward type (money, chocolate milk) and timepoint (run 1, run 2). Results revealed a significant main effect of reward type for both ACC (F (1, 31) = 5.45, P < 0.05, partial η^2^ = 0.15) and RT (F (1, 31) = 6.72, P < 0.05, partial η^2^ = 0.18), suggesting higher ACC and faster responses for money (P < 0.05), albeit no significant interaction effect between the factors was observed, suggesting that monetary and chocolate milk had overall different incentive values (**Figure 3C**)

### 3.4 fNIRS results

#### 3.4.1 Anticipation phase

Examining neural activity during the anticipation phase in experiment 2 using a repeated ANOVA with the factors reward condition (2: chocolate milk, money), time(2: run 1, run 2)and channel(1-19), revealed a significant main effect of channel (F (18, 588) = 3.24, P < 0.01, partial η^2^ = 0.10), a significant reward x channel interaction effect (F (18, 588) = 4.67, P < 0.01, partial η^2^ = 0.13) as well as a significant time x channel interaction effect (F(18, 588) = 2.50, P < 0.05, partial η^2^ = 0.08). Channel-specific post-hoc t-tests revealed that vlPFC/lOFC activation, particularly in the right hemisphere, increased during the anticipation of money over the course of the experiment (money_run1 vs. money_run2: channel 1, P _FDR_ = 0.03; channel 2, P _FDR_ = 0.02; channel 3, P _FDR_ = 0.02; channel 4, P _FDR_ = 0.03; channel 19, P _FDR_ = 0.04). Comparing the reward-types separately during the runs, further revealed that vlPFC/lOFC regions exhibited stronger activation for money as compared to chocolate milk during the second run albeit not for the first run (money_run2 vs. chocolate milk_run2: channel 1, P _FDR_ < 0.001; channel 2, P _FDR_ = 0.05; channel 3, P _FDR_ = 0.01; channel 18, P _FDR_ = 0.01; channel 19, P _FDR_ = 0.05; all p-values in run 1 > 0.04; **Figure 4A**). Together, these findings suggest that the vlPFC/lOFC tracks the changes of value, specifically the increasing motivation for the monetary rewards.

**Figure 4.**
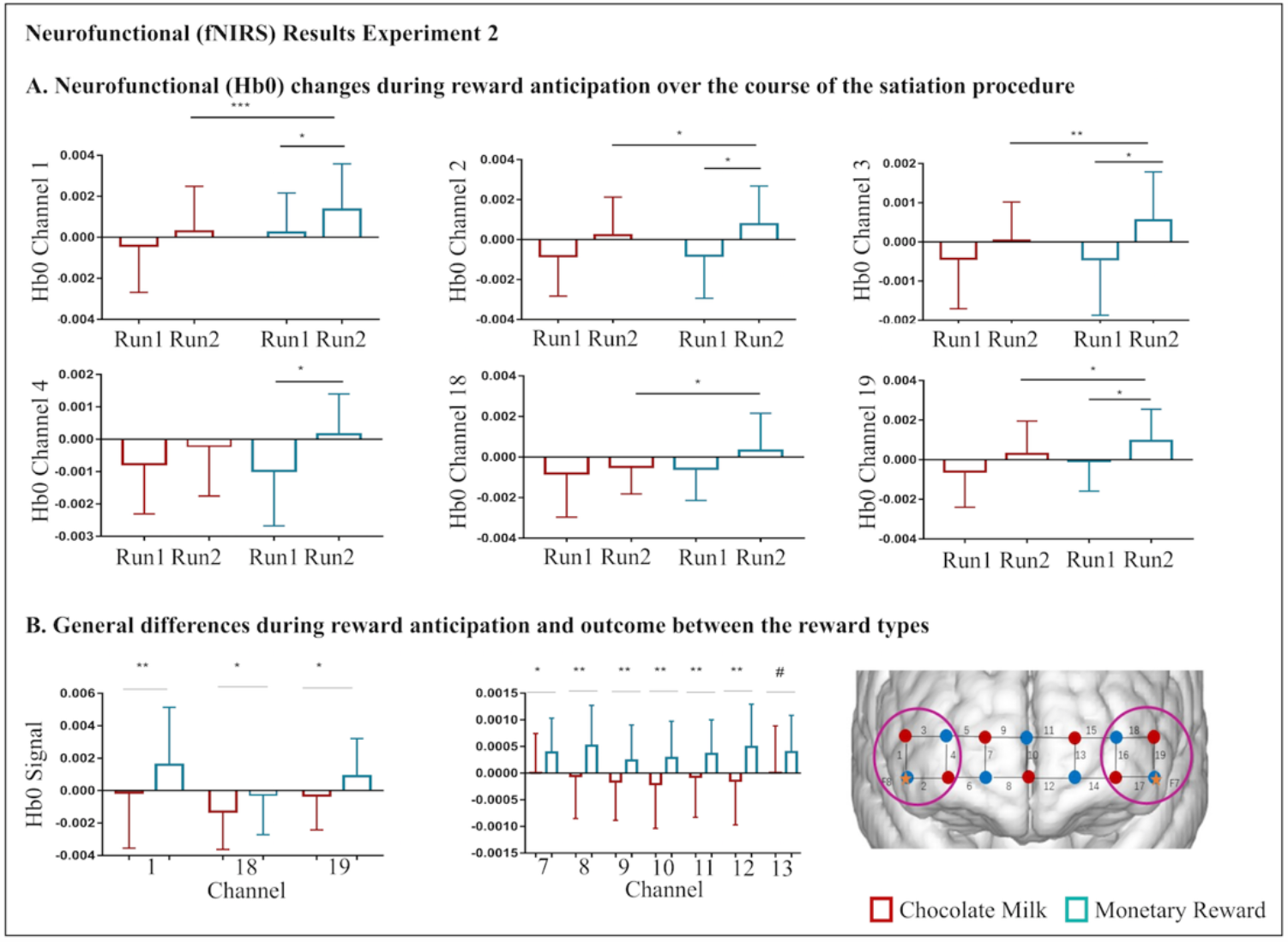
Neurofunctional (fNIRS) results in Experiment 2. **(A)** Examination of changes between the first and the second run of the incentive delay task revealed that activation in lateral prefrontal regions (channels 1,2,3,4,19) during the anticipation of money increased over the course of the experiment. These regions (channels 1,2,3,18,19) moreover exhibited stronger activation during the anticipation of money as compared to chocolate milk during the second run. **(B)** Examination of general activation differences revealed that lateral prefrontal channels (1, 18, 19) exhibited generally higher activity during the anticipation of money as compared to chocolate milk (left panel). During the outcome phase medial prefrontal regions (channels 7-12) exhibited generally higher activity for money as compared to chocolate milk (middle panel). Location of fNIRS optodes: lateral parts (8 channels covering channels 1-4 right and channels 16-19 left) of PFC); medial parts (11 channels, channels 5-15) of the PFC # p _FDR_*=0.06, * P _FDR_ < 0.05, ** P_FDR_ < 0.01.

In an additional analysis we omitted the factor ‘run’ to explore overall differences in the neural coding of primary and secondary rewards. Combining the runs in a repeated ANOVA with condition (2: chocolate milk, money) × channel (1-19) revealed a main effect of channel (F (18, 558) = 3.24, P < 0.01, partial η^2^ = 0.10) and condition x channel interaction effect (F (18, 588) = 4.67, P < 0.01, partial η^2^ = 0.13). Post-hoc comparisons demonstrated that the most anterior parts of the vlPFC/lOFC exhibited higher brain activity during the anticipation of secondary (money) as compared to primary (chocolate milk) rewards (channel 1, P _FDR_ = 0.003; channel 18, P _FDR_ = 0.04; channel 19; P _FDR_ = 0.02) (**Figure 4B, left panel**).

#### 3.4.1 Outcome phase

Examining neural activation during the outcome phase by means of a repeat ANOVA with the factors condition (2: chocolate milk, money), time (2: run 1, run 2)and channel (1-19) revealed a significant main effect of reward (F (1, 31) = 4.36, P < 0.05, partial η^2^ = 0.12) and a channel x reward interaction effect (F (18, 588) = 7.09, P < 0.001, partial η^2^ = 0.19). For the outcome phase no significant main or interaction effects of time were observed, suggesting that outcome-related neural activation did not change over the experiment.

To further explore differences in the neural coding of primary and secondary (monetary) rewards, the data was again pooled over the runs. Repeated ANOVAs with reward condition (2: chocolate milk, money) and channel (1-19) revealed significant main effects of condition (F (1, 31) = 4.36, P < 0.05, partial η^2^ = 0.12) and a channel x condition interaction (F (18, 588) = 7.09, P < 0.001, partial η^2^ = 0.19). Post hoc T tests revealed that monetary outcomes induced significantly higher neural activity compared to chocolate milk in the vmPFC/mOFC (channel 7-12, all P _FDR_ < 0.05) (**Figure 4B, middle panel**).

## 4. Discussion

The present study aimed at determining differences in the devaluation of primary (chocolate milk) and secondary (money) rewards on the behavioral and neural level. Consistent with previous research, the value of the primary reward decreased over the course of the satiation procedure. Although the hedonic value of the primary and secondary rewards did not differ at the beginning of the satiation procedure, the hedonic evaluation of the secondary reward increased over the experiments such that money was rated as more pleasant than chocolate milk after the satiation procedures. Opposite devaluation trajectories for the primary and secondary reward were also reflected in the behavioral indices acquired during the satiation procedure. Choice preference and decision time did not as yet differ at early stages of the experiment. Over the course of the experiment both indices changed in the expected direction for the primary reward. In contrast, preference for money increased and corresponding decision times decreased, leading to significantly less as well as slower choices of the natural reward (Experiment 1). In Experiment 2, participants exhibited an overall higher accuracy and faster reaction times for the secondary as compared to the primary reward, suggesting an overall higher motivation to perform the incentive delay paradigm when money was at stake. On the neural level, -irrespective of satiation-dependent changes - anticipation of the secondary reward was associated with higher activation in the anterior VLPFC/lOFC, whereas it was associated with higher activity in the most anterior part of the VMPFC/mOFC during the outcome phase. Interestingly, VLPFC/lOFC anticipatory, and not outcome-related activation, increased for secondary rewards over the satiation procedure, whereas no changes were observed for the primary reward. Importantly, during early stages of the procedure anticipatory VLPFC/lOFC activation did not differ by reward conditions. Notably, in contrast during later stages, the VLPFC/lOFC exhibited stronger anticipatory activation for secondary as compared to primary rewards, suggesting that these changes are the neural representations of the increasing value of the secondary reward.

Consistent with several previous studies that examined sensory-specific satiation of natural rewards with repeated exposure or consumption, we observed a strong devaluation of the primary reward as reflected by decreased hedonic value and choice preference as well as increasing decision times [9–12]. Conversely, a very different pattern was observed for the secondary reward, such that hedonic value and choice preference increased while decision times decreased. Devaluation effects in satiety procedures employing primary rewards are considered a decrease in pleasure and reward value, which are regulated by adaptations in the homeostatic and hedonic state to facilitate appropriate goal-directed behavior. Secondary rewards are not directly coupled to innate homeostatic balance or hedonic satiety, and while both primary and secondary rewards engage key regions of the brain’s reward system, the lack of regulatory devaluation mechanisms for secondary rewards may lead to progressively increasing reward value with repeated exposure to secondary rewards, especially with money or opioids.

A similar process – albeit on a considerably longer timeframe – has been observed for drugs of abuse, which are highly potent secondary reinforcers. Previous studies suggest that the anticipation and receipt of monetary rewards in healthy individuals and exposure to drug-associated cues in recreational drug users elicit strong activity in the striato-prefrontal reward circuits of the brain[54, 55]. Rodent models have provided compelling evidence for escalating drug use by employing extended drug-self administration procedures suggesting that a progressive increase in the hedonic set point of the individual [56] and in the incentive salience of the secondary reward stimulus[57], may drive escalating use during early stages of the addictive process. Whereas these drug-use models observed changes over the course of days, we observed a similar, yet more subtle, pattern during the repeated receipt of small amounts of monetary reinforcers within a single session. The strong motivation to obtain money has been conceptualized in different frameworks, with some recent theories proposing biological and conceptual similarities with drug addiction having recently gained increasing support [58]). The anticipation and receipt of primary as well as secondary rewards including drugs as well as money has been associated with increasing dopamine (DA) release in the striatum, particularly the ventral striatum[59, 60] which exhibits strong connections with the OFC[61]. Stronger stimulant drug-induced DA release in the ventral striatum is associated with stronger subjective hedonic experience of drug use [62, 63] and increases goal-directed behavior towards monetary rewards despite increasing efforts necessary to obtain these [64]. Although a clinical picture of ‘money addiction’ has not been described it has been proposed as a driving factor behind some behavioral addictions such as pathological gambling [58] which – during its early stages – is characterized by the motivation of gaining more money [65].

Whereas the behavioral assessments did not allow us to determine differential effects during reward anticipation (‘wanting’, incentive motivation) versus during reward receipt (‘liking’, pleasure) the concomitant fNIRS acquisition during the incentive reward experiment was more informative. The secondary reward was associated with overall higher activity in the anterior lateral part of the OFC during reward anticipation and higher activity in the most medial OFC during reward outcome. Differential changes over the satiation procedure specifically occurred in the lateral OFC during the anticipation phase of the secondary reward. In the satiation period higher medial OFC activation during the receipt of secondary rewards aligns with findings from a previous meta-analysis of fMRI studies suggesting that monetary outcomes specifically engage the most anterior portion of the medial OFC [17]. Moreover, previous studies suggested that medial OFC activation during reward consummation is positively associated with subjective pleasantness (liking) [66–70] suggesting that the medial OFC codes reward magnitude during consumption [71] and overall outcome value to guide goal-directed behavior [11, 20, 72]. The present findings make ‘biological sense’ that abstract secondary rewards are encoded in evolutionarily more recent brain regions[18], or alternatively reflect a generally higher salience of the monetary as compared to the food reward. The latter interpretation is additionally supported by generally faster response times and higher accuracies for monetary rewards during the incentive delay task.

Whereas the medial OFC encodes general economic value, the lateral OFC serves more complex functions and is thought to represent the current value of a reward. This is shown by lesions of this region that impair an animal’s ability to assign reward value to a specific choice [27], which in turn hampers the ability to adapt choice behavior during devaluation procedures [73]. Moreover, the lateral OFC has been associated with the dynamic updating of reward type-specific value [26, 74] by way of integrating prior with current information [75]. The lateral OFC also encodes reward-type specific representations which are modulated by selective devaluation to primary rewards such as food odor[11]. Generally higher activity for the secondary reward in combination with increasing anticipatory activity in this region over the devaluation procedure likely reflects an overall stronger identity-specific coding of monetary rewards as well as increasing incentive motivation of the monetary reward with repeated receipt of the outcome.

Contrary to previous fMRI studies, in our experiment we did not observe neural changes that accompanied the devaluation of the primary reward over the satiation procedure. This may be partly explained by the limited acquisition depth of fNIRS. Previous studies reported food satiation associated decreases during the expectation phase in regions such as the ventral striatum (VS), supramarginal gyrus (SMG), insula or amygdala [20, 29, 76], which were not covered by the present fNIRS acquisition protocol. In addition to limitations inherent to the neuroimaging technology other limitations of the present study include the focus on young male subjects with normal weight. Given that previous studies demonstrated decreased sensitivity to reward devaluation in obese individuals[77], gender differences in reward responses[78] and age-related changes during anticipation including receipt of monetary rewards [79], the generalization of the present findings remains to be established in future studies.

In summary, our findings demonstrate for the first time that primary and secondary rewards show opposite devaluation processes over a satiation procedure such that the value of chocolate milk decreased whereas the value of money increased. On the neural level, increases in the value of the secondary reward were paralleled by increasing anticipatory activity in the lateral OFC, suggesting an identity-specific increased incentive motivation to obtain the secondary reward with repeated exposure.

## Acknowledgements

This work was supported by the National Key Research and Development Program of China (Grant No. 2018YFA0701400), National Natural Science Foundation of China (NSFC No. 31700998, 31530032).

